# IKKalpha-Mediated Non-canonical NF-kappaB Signaling is Required to Support Murine Gammaherpesvirus 68 Latency *In Vivo*

**DOI:** 10.1101/2022.01.05.475165

**Authors:** Brandon Cieniewicz, Varvara Kirillov, Isabel Daher, Xiaofan Li, Darby G. Oldenburg, Qiwen Dong, Julie A. Bettke, Kenneth B. Marcu, Laurie T. Krug

**Author notes:** Duchossois Family Institute, University of Chicago, Chicago, IL, USA.

## Abstract

Non-canonical NF-kappaB signaling is activated in B cells via TNF receptor superfamily members CD40, Lymphotoxin beta-R, and BAFF-R. The non-canonical pathway is required at multiple stages of B-cell maturation and differentiation, including the germinal center reaction. However, the role of this pathway in gammaherpesvirus latency is not well understood. Murine gammaherpesvirus 68 (MHV68) is a genetically tractable system used to define pathogenic determinants. Mice lacking the BAFF-R exhibit defects in splenic follicle formation and are greatly reduced for MHV68 latency. We report a novel approach to disrupt non-canonical NF-kappaB signaling exclusively in cells infected with MHV68. We engineered a recombinant virus that expresses a dominant negative form of IKKalpha, named IKKα-SA, with S176A and S180A mutations that prevent phosphorylation by NIK. We controlled for the transgene insertion by introducing two all-frame stop codons into the IKKα-SA gene. The IKKα-SA mutant but not the IKKα-SA.STOP control virus impaired LTbetaR-mediated activation of NF-kappaB p52 upon fibroblast infection. IKKα-SA expression did not impact replication in primary fibroblasts or in the lungs of mice following intranasal inoculation. However, the IKKα-SA mutant was severely defective in colonization of the spleen and in the establishment of latency compared to the IKKα-SA.STOP control and WT MHV68 at 16 dpi. Reactivation was undetectable in splenocytes infected with the IKKα-SA mutant, but reactivation in peritoneal cells was not impacted by IKKα-SA. Taken together, the non-canonical NF-kappaB signaling pathway is essential for the establishment of latency in the secondary lymphoid organs of mice infected with the murine gammaherpesvirus pathogen MHV68.

**IMPORTANCE:** The latency programs of the human gammaherpesviruses EBV and KSHV are associated with B cell lymphomas. It is critical to understand the signaling pathways that are used by gammaherpesviruses to establish and maintain latency in primary B cells. We used a novel approach to block non-canonical NF-kappaB signaling only in the infected cells of mice. We generated a recombinant virus that expresses a dominant negative mutant of IKKalpha that is non-responsive to upstream activation. Latency was reduced in a route- and cell type-dependent manner in mice infected with this recombinant virus. These findings identify a significant role for the non-canonical NF-kappaB signaling pathway that might provide a novel target to prevent latent infection of B cells with oncogenic gammaherpesviruses.

## INTRODUCTION

Herpesviruses use a strategy of latency to achieve lifelong persistence in the host. While gammaherpesviruses infect and grow productively in multiple cell types including epithelial cells, endothelial cells, and macrophages, B cells are the predominant reservoir of latency, defined by a non-integrated viral genome with highly circumscribed viral gene expression to evade detection by the immune system. In the context of host immunosuppression, the control of viral latency and the suppression of productive infection is impaired, predisposing the host to the development of malignancies (1). The human gamma-herpesviruses Epstein-Barr virus (EBV, HHV-4) and Kaposi Sarcoma herpesvirus (KSHV, HHV-8) have etiological associations with multiple lymphomas and malignancies (2-9). Despite the importance of latency in the viral life cycle, the mechanisms that drive latency establishment and the cellular signals that maintain latency or initiate reactivation are poorly understood, hindering the development of therapeutics against infection and virus-associated malignancies.

Murine gammaherpesvirus 68 (MHV68, MuHV-4, γHV68) is genetically colinear to KSHV, and provides a tractable model to discover and understand the host signaling pathways that contribute to the establishment and maintenance of a latent gammaherpesvirus infection (10). Infection of mice leads to a chronic infection involving B cell and macrophage latency which can manifest lymphoproliferative disease if immune control is impaired. MHV68 infection drives infected naïve B cells to enter the germinal center, undergo proliferative expansion, and ultimately establish viral latency in long lifespan memory B cells (10).

The NF-kappaB (NF-κB) signaling pathway is well known for its roles in cell survival, proliferation, and the induction of inflammatory cytokine responses. NF-κB also plays a critical role in multiple steps of B cell development and activation, and germinal center-dependent B cell responses including class switch recombination (11). NF-κB signaling occurs via two distinct pathways, the canonical and non-canonical NF-κB signaling pathways. Canonical NF-κB signaling activates the transcription factors p105/p50, RelA and c-Rel in numerous cell types through cytokine receptors, toll-like receptors, and antigen receptors specifically in lymphocytes. Activation of this pathway leads to phosphorylation of the inhibitory molecule IκBα, leading to its rapid degradation and release of transcription factors to mediate signaling. Non-canonical NF-κB signaling is activated in a more restricted manner through the stimulation of multiple receptors belonging to Tumor Necrosis Factor (TNF) super family, including CD40 (12), B cell activating factor (BAFF) (13) and lymphotoxin-β (LTβ) (14). Non-canonical NF-κB signaling is characterized by slow kinetics and a dependence on protein synthesis (15, 16). NF-κB-inducing kinase (NIK), the critical mediator of non-canonical NF-κB signaling, is constitutively degraded by TRAF2 and TRAF3. Activation of the non-canonical signaling pathway leads to the phosphorylation and degradation of TRAF3, allowing NIK to slowly accumulate through *de novo* protein synthesis. NIK accumulation leads to phosphorylation of IKKalpha (IKKα), which in turn phosphorylates the p100 transcription factor, causing ubiquitin-mediated processing of p100 to the mature NF-κB p52 subunit and the subsequent nuclear translocation of RelB/p52 heterodimers (17).

Infection of B cells by the human gammaherpesviruses leads to constitutive NF-κB activation through the expression of viral proteins and miRNAs that mimic endogenous receptor signaling (18-20). While these molecules are important for infected cell survival *in vitro*, the role of both endogenous and virus-driven NF-κB signaling *in vivo* is not well-defined. Previous work by our laboratory examined the role of the canonical NF-κB signaling pathway in latency establishment and found that expression of a dominant negative inhibitor of canonical signaling reduced viral latency and reactivation in B cells (21). In addition, p50^-/-^ mice infected with MHV68 were characterized by persistent replication in the lungs and heightened latency in the spleen and a lack of virus-specific antibody production (22). This broad defect in control of replication led us to generate mixed (WT/p50^-/-^) bone marrow chimeric mice wherein the WT hematopoietic-derived cell population would restore immune control. This competitive model of infection revealed a tremendous skew in the population of cells that supported MHV68 latency. B cells lacking p50 had a sustained and substantial defect in the frequency of viral genome-positive cells compared to their WT counterparts three months after infection.

Non-canonical NF-κB signaling is similarly critical for the development, maturation, and survival of B cells. Deletion of the CHUK/*nfκb2* gene, which encodes the IKKα protein (23), results in defects in B cell function and deformed microarchitecture in peripheral lymphoid organs (24). Alymphoplasia (aly) mice carrying a loss of function mutation in NIK are characterized by the systemic absence of lymph nodes and disorganized splenic and thymic structures with severe immunodeficiency (25). Mice lacking the BAFF-R exhibit defects in splenic follicle formation and are greatly reduced for MHV68 latency (26). Non-canonical NF-κB signaling is seemingly central to the cellular reservoirs in which gammaherpesviruses establish latency, but its direct role in facilitating latency is difficult to ascertain given the broad loss of follicular architecture in knock-out mice.

Here, we sought to examine the role of IKKα-mediated non-canonical NF-κB signaling in latency establishment using an approach that would specifically impair IKKα function in the cells infected with MHV68. We generated a virus that expresses a dominant negative IKKα molecule that impairs non-canonical NF-κB signaling to elucidate the contribution of IKKα in MHV68 replication and latency establishment *in vivo*. We found that lytic replication was not affected upon introduction of dominant negative IKKα in cell culture or in mice. However, establishment of viral latency in the spleen after intranasal inoculation was reduced to a level below our limit of detection. Interestingly, latency establishment in the peritoneal compartment was unimpaired, although establishment of latency in the splenic reservoir after intraperitoneal inoculation was significantly impaired. These are the first data to report an intrinsic requirement for NIK-dependent non-canonical NF-κB signaling in the establishment of gammaherpesvirus latency in the splenic compartment of the host.

## RESULTS

### Generation of recombinant viruses expressing dominant-negative IKKα-SA

Processing of p100 is triggered upon induced phosphorylation at the carboxyl-terminus of p100, which is regarded as the central event in activation of the non-canonical NF-κB pathway. The phosphorylation of p100 by IKKα is licensed by the phosphorylation of two serines in its activation loop by the upstream kinase NIK. We previously employed transgene expression of the transdominant IκBαM to suppress NF-κB signaling in the infected B cells of mice (21). To specifically impair non-canonical NF-κB signaling in the infected cell, we engineered a recombinant MHV68 expressing a dominant negative form of IKKα, IKKα (S176A,S180A) (IKKα-SA) (27), from a neutral intergenic locus between ORF57 and ORF58 in the MHV68 genome (28). IKKα-SA bears two serine to alanine mutations in the T loop that prevents phosphorylation by NIK and downstream signaling (29-31). The IKKα-SA ORF was followed by an internal ribosome entry sequence (IRES) and a histone 2B-YFP reporter gene **(FIG 1A)**. Two independent clones of this recombinant MHV68, termed IKKα-SA1 and -SA2, were generated.

**FIG 1.**
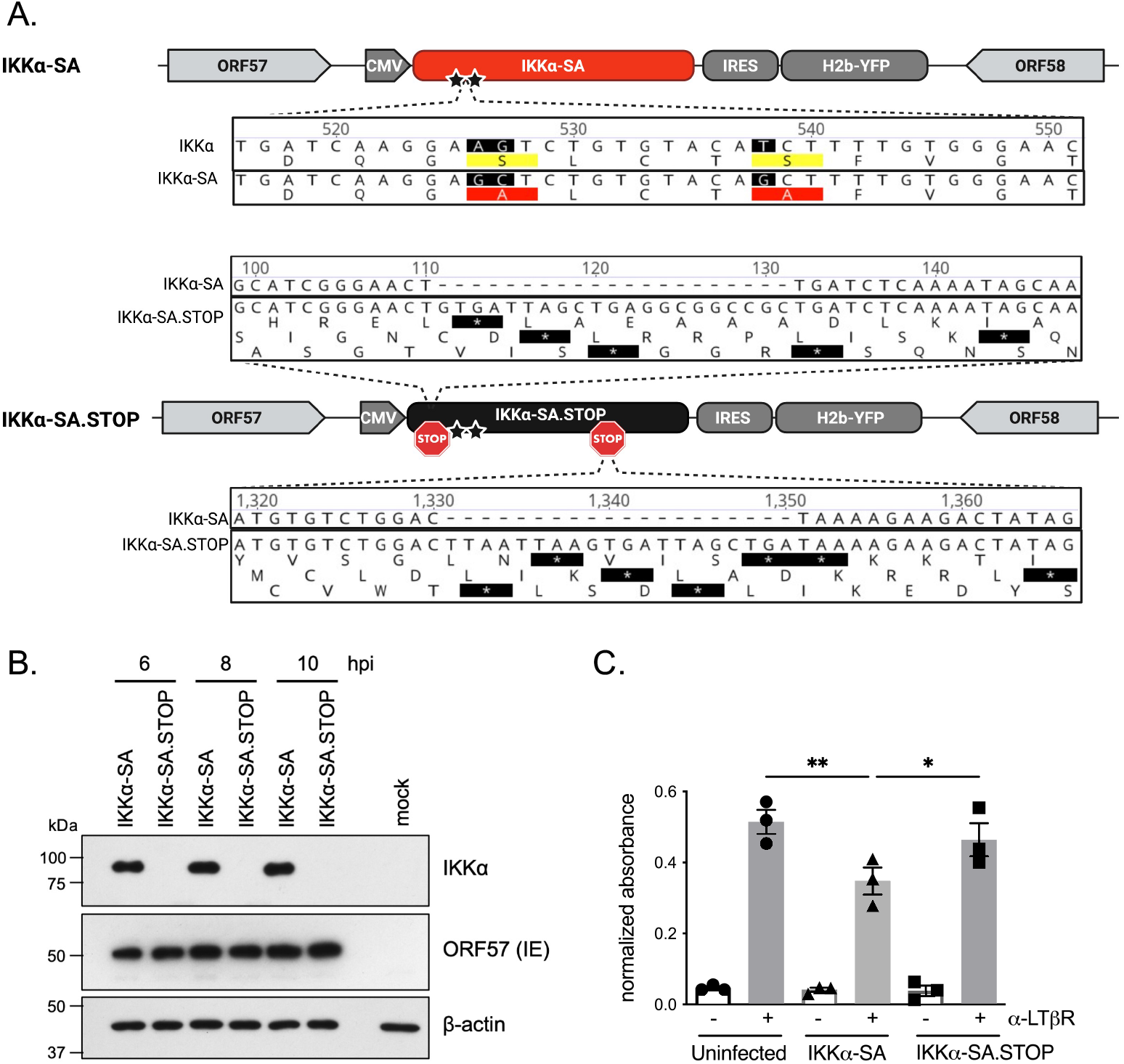
Generation of recombinant MHV68 expressing transgene that impairs IKKα signaling. (A) Schematic of IKKα-SA and IKKα-SA.STOP viruses made with Biorender. A cassette encoding a CMV-IE promoter driving IKKα-SA or IKKα-SA.STOP, an IRES, and an H2b-YFP was inserted into the neutral locus between ORF57 and ORF58 using BAC-mediated recombination. Stars denote location of S176A and S180A mutations; STOP indicates all-frames stop cassette insertion. (B) Primary WT MEFs were infected with IKKα-SA or IKKα-SA.STOP at an MOI of 10. Immunoblots of protein lysates were analyzed at 6, 8, and 10 hpi for the IKKα transgene and the IE viral protein ORF57. β-actin was used as a loading control. (C) Primary MEFs were infected with MHV68-IKKα-SA or IKKα-SA.STOP at an MOI of 10. At 4 hpi, cells were stimulated with 0.3 μg/ml of αLTβR antibody for 5 h. At 9 hpi, cells were harvested. Nuclear fractions were probed for p52 activation using a p52 ELISA and normalized to uninfected, unstimulated cells. Results represent mean +/- SEM for three independent experiments with duplicate samples per replicate. ***, p<0.001; ****, p<0.0001; analyzed using ANOVA with Sidak’s multiple comparison post-test.

To control for changes in viral fitness due to gene insertion into the ORF57-58 locus, we also generated a virus bearing two all-frames stop codon sequences after the start codon of IKKα and prior to the gene region encoding the T-loop containing the mutated serine residues (29). The recombinant viruses were generated using BAC-mediated *en passant* recombination into a previously sequence-verified version of the IKKα-SA BAC **(FIG 1A)**. Two independent clones of this recombinant MHV68, termed IKKα-SA.STOP1 and -SA.STOP2, were generated. Purified viral BACs were analyzed using restriction fragment length polymorphism **(FIG S1)** and whole genome sequencing to verify that the intended mutations were present and that secondary mutations were absent. As designed, when compared to NM_001278 (hIKKα CDS), the IKKα-SA and IKKα-SA.STOP genomes had AG to GC substitutions at nt 526-527 and a T to G substitution at nt 538 to generate S176A and S180A amino acid changes in the IKKα CDS, respectively. The IKKα-SA.STOP genome had the expected insertions at nt 111 and 1310 that led to a translational stop in the IKKα-SA ORF. Additional mutations were uncovered in the IKKα-SA ORF for both IKKα-SA and IKKα-SA.STOP viruses: a G to A change at nt 802 of IKKα.SA that leads to a non-synonymous V to I coding change; and a synonymous C to T change nt 1,009.

To confirm transgene expression from the mutant viruses, murine embryonic fibroblasts (MEFs) were infected with the IKKα-SA and IKKα-SA.STOP viruses at an MOI of 5 and lysates were collected 6, 8, and 10 hours post infection (hpi). IKKα was expressed at high levels in cells infected with the IKKα-SA virus, but not in cells infected with the IKKα-SA.STOP **(FIG 1B)**.

We next tested for impairment of non-canonical NF-κB pathway signaling by infection with the MHV68 recombinant expressing IKKα-SA. Activating the lymphotoxin beta receptor (LTβR) in MEFs by a LTβR antibody induces non-canonical signaling (32). MEFs were stimulated for 18 h with antibody against the LTβR and tested for p100 cleavage in the whole cell, cytoplasmic, and nuclear fractions. We also stimulated MEFs with the canonical NF-κB pathway activator TNFα for 15 min. NF-κB p100 cleavage and p52 translocation was observed only after αLTβR stimulation **(FIG S2)**. Next MEFs were infected with IKKα-SA and IKKα-SA.STOP virus at an MOI of 10 for 4 h prior to stimulation of infected cells with αLTβR antibody for 5 h. The levels of activated p52 in nuclear fractions that recognized immobilized oligonucleotides with p52 consensus binding sites were measured in a p52 NF-κB ELISA. NF-κB p52 binding activity in nuclear fractions was induced upon stimulation with αLTβR antibody in the uninfected cultures **(FIG 1C)**. Infection with IKKα-SA led to a significant reduction of p52 binding that was not observed in cells infected with IKKα-SA.STOP. The reduction in nuclear p52 in infected, stimulated cells indicates that the recombinant MHV68 expressing IKKα-SA impaired non-canonical NF-κB signaling infection.

### IKKα-SA does not impair lytic replication

Next, we assessed if transgene overexpression impaired MHV68 replication in primary fibroblasts. MEFs were infected with IKKα-SA1 and -SA2, IKKα-SA.STOP1 and .STOP2, or WT MHV68 at a low multiplicity of infection (MOI 0.05) to allow for multiple rounds of replication. The IKKα-SA and IKKα-SA.STOP viruses replicated with similar kinetics in a multistep growth curve **(FIG 2A)**, indicating that IKKα-SA expression does not impair lytic replication. We also investigated the impact of the loss of non-canonical NF-κB signaling by infecting MEFs generated from *CreER*^*T2*^*/IKKα*^*fl/fl*^ mice that are inducible for IKKα exon deletion **(FIG S3A)**. WT MHV68 replication was not altered in MEFs knocked out for IKKα **(FIG S3B)**. Replication in the lung tissue provides a stringent analysis of factors that influence productive infection. Intranasal infection of C57BL/6 mice with 1,000 PFU of IKKα-SA or IKKα-SA.STOP viruses resulted in similar levels of replication in the lungs of infected mice as compared to WT virus infection at 4 and 9 days post-infection (dpi) **(FIG 2B)**. Taken together, the loss of IKKα-mediated non-canonical NF-κB signaling does not impair virus replication.

**FIG 2.**
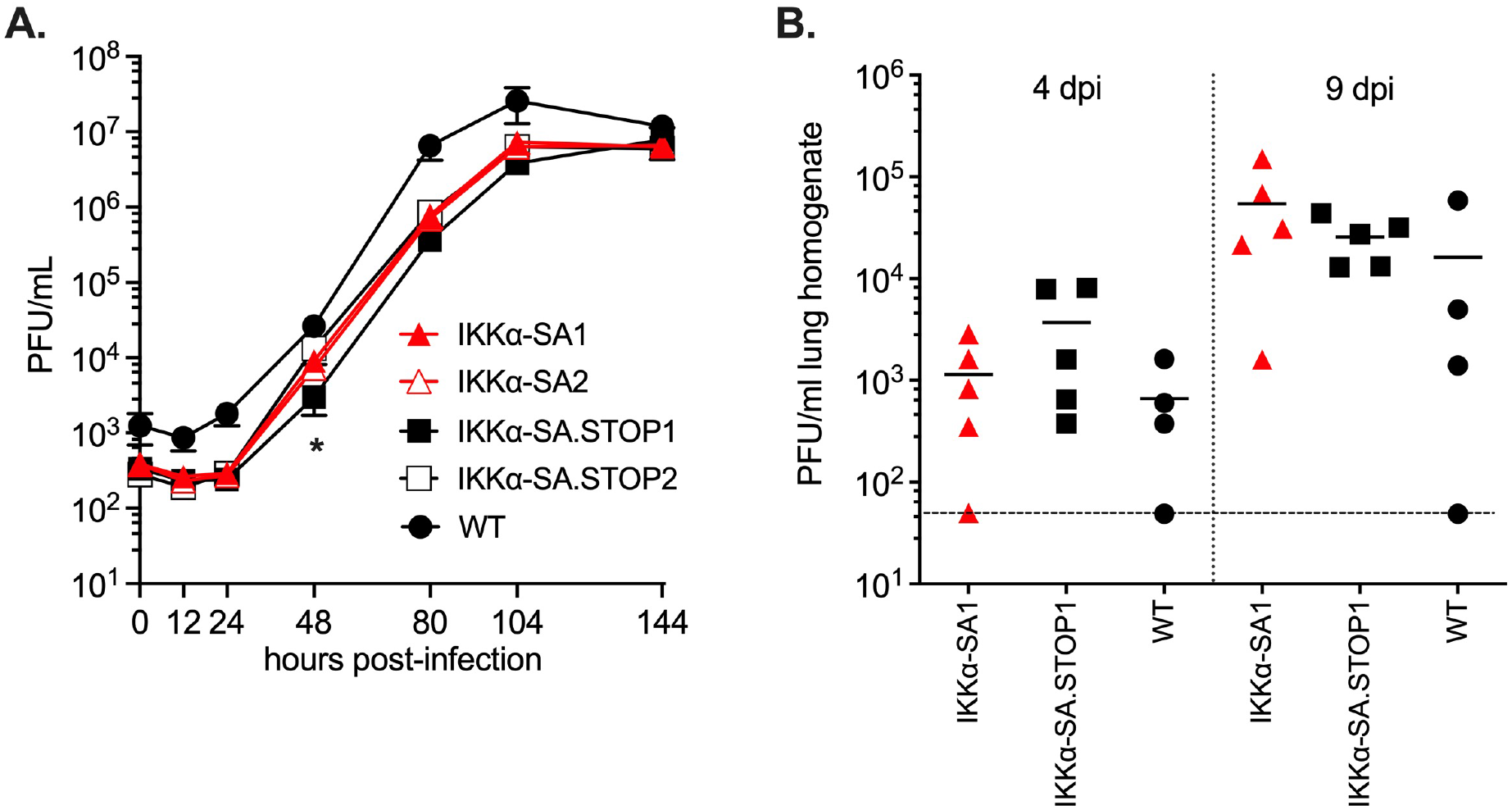
Lytic replication is not altered by the expression of IKKαSA in murine fibroblast cells or in the lungs of infected mice. (A) Murine embryonic fibroblasts infected at MOI = 0.05 and cell lysate harvested at the indicated time points. Viral titer determined by plaque assay and measured in triplicate. Results analyzed using two-way ANOVA with Tukey’s multiple comparisons. *, p<0.05 at 48 hpi; WT MHV68 differs from other viruses at all time points (p <0.05) due to input difference. (B) C57BL/6 mice were infected with 1,000 PFU of the indicated viruses via the intranasal route. Lungs were removed and homogenized at the indicated times post-infection, and virus titers were determined by plaque assay. Symbols represent titer per ml lung homogenate for individual mice, lines represent the mean titers; dashed line indicates limit of detection (50 PFU/ml). ‘WT’ indicates BAC-derived MHV68. No significant differences found by one-way ANOVA for each timepoint.

### IKKα-mediated non-canonical NF-κB activation is a critical host signaling pathway for splenic latency

Given that NIK-mediated non-canonical NF-κB signaling occurs upon engagement of many cytokines and surface molecules of cells that engage B cells in the lymphoid tissue, we hypothesized that interference with this pathway via expression of the IKKα-SA would have a negative impact on latency in splenocytes. After intranasal infection, MHV68 disseminates to the spleen via B cells, reaching the peak of latency ∼16 dpi. An initial indicator of this colonization event is splenomegaly. Sixteen days after intranasal infection with 1000 PFU, WT infection led to a two-fold increase in spleen weight as compared to naïve mice **(FIG 3A)**. Infection with the IKKα-SA viruses led to a significant loss in splenomegaly compared to the control IKKα-SA.STOP or WT MHV68 infection. Limiting dilution PCR is a quantitative measurement of the frequency of splenocytes that harbor the viral genome and reflects the establishment of latency. Infection with the IKKα-SA virus led to a significant two-log reduction in genome-positive splenocytes (1/27,295 splenocytes) compared to infection with WT virus (1/227) or with the IKKα-SA.STOP control (1/393) **(FIG 3B)**.

**FIG 3.**
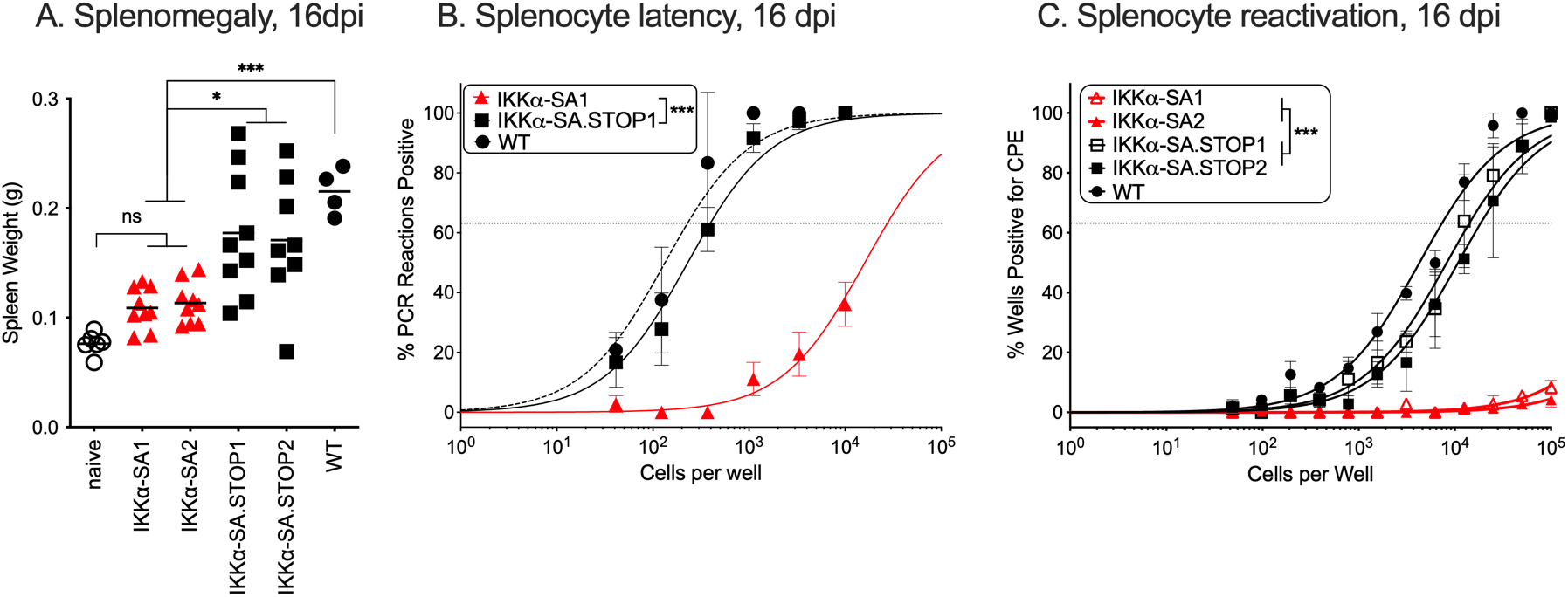
Latency establishment in the spleen is severely ablated at 16 dpi by the expression of IKKα-SA. C57BL/6 mice were infected with 1,000 PFU of the indicated viruses via the intranasal route and tissues were analyzed 16 dpi. (A) Spleens weights from individual uninfected, naïve or infected mice are represented by symbols and lines indicate mean weight. Data from two independent experiments of 3-5 infected mice per group. Significance was determined by one-way ANOVA with Tukey’s multiple comparison test; *, p<0.05; ***, p<0.0009. (B) Latency was measured in bulk splenocytes by determining the frequency of viral genome-positive splenocytes by limiting dilution PCR. Briefly, intact splenocytes were serially diluted and subjected to nested PCR reactions to detect viral ORF50. (C) Reactivation was measured by coculturing serial dilutions of intact splenocytes from infected mice onto a monolayer of MEFs and scoring for CPE 2-3 weeks post-coculture. For the limiting dilution analyses, curve fit lines were determined by nonlinear regression analysis. Using Poisson analysis, the intersection of the nonlinear regression curves with the dashed line at 63.2% was used to determine the frequency of cells that were either positive for the viral genome (B) or reactivation (C). Data are generated from three independent experiments with 3 to 5 mice per group for IKKα-SA and IKKα-SA.STOP viruses and two independent experiments with WT BAC-derived MHV68. Error bars indicate SEM. Significance was determined by unpaired, two-tailed t-test (B) ***, p<.001 or one-way ANOVA with Tukey’s multiple comparisons (C) ***, p<0.0001.

Reactivation from latency was analyzed using a limiting dilution *ex vivo* coculture assay. Splenocytes from mice infected with WT virus (1/7,292) or either of the IKKα-SA.STOP control viruses (1/14,033 for .STOP1 and 1/18,374 for .STOP2) had comparable frequencies of spontaneous reactivation upon explant **(FIG 3C)**. Consistent with the severe defect in the establishment of latency observed in FIG 3B, there was a near complete ablation of cytopathic effect in MEFs cocultured with 100,000 splenocytes from mice infected with IKKα-SA viruses, well below the limit of accurate enumeration **(FIG 3C)**. Thus, while there was no defect in acute replication in the lung tissue after intranasal inoculation, interference with NIK-mediated non-canonical NF-κB signaling via IKKα-SA expression severely curtailed the establishment of latency in splenocytes.

A more permissive route of MHV68 infection is intraperitoneal administration. Intraperitoneal infection of C57BL/6 mice with 1000 PFU of IKKα-SA.STOP1 i.p. led to a mean three-fold increase in splenomegaly over uninfected mice **(FIG 4A)**. There was a significant diminishment in the weight of spleens in mice infected with IKKα-SA1. Latency establishment was reduced five-fold in the splenocytes of mice infected with IKKα-SA1 (1/1,018) compared to mice infected with IKKα-SA.STOP1 (1/214) **(FIG 4B)**. A similar six-fold reduction in reactivation of latency manifested in the splenocytes of mice infected with IKKα-SA1 (1/70,733) compared to mice infected with IKKα-SA.STOP1 (1/11,159) **(FIG 4C)**. Long-term latency in splenocytes at 42 dpi was nearly comparable between mice infected with IKKα-SA1 (1/9,942) or IKKα-SA.STOP1 (1/3,837) **(FIG 4D)**. Altogether, latency and the ensuing reactivation in splenocytes is impaired with the expression of the inhibitory IKKα mutant in the infected cells.

**FIG 4.**
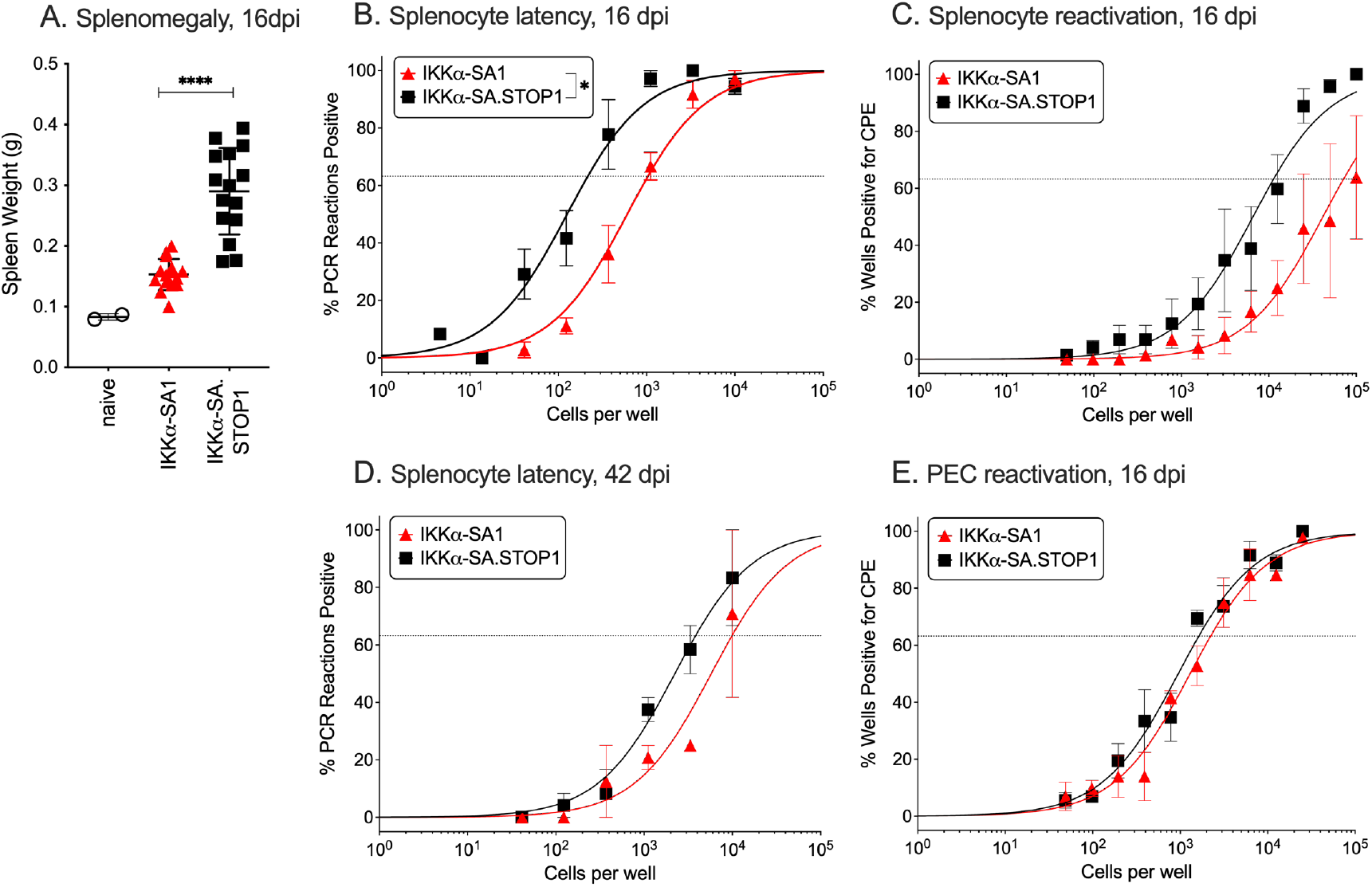
Latency establishment in the spleen, but not the peritoneal compartment, is impaired by the expression of IKKα-SA. C57BL/6 WT mice were infected with 1,000 PFU of the indicated viruses via the intraperitoneal route. (A) Spleens weights from individual uninfected, naïve or infected mice are represented by symbols and lines indicate mean weight. Data from three independent experiments of 5 infected mice per group. Significance was determined by unpaired two-tailed t-test; ****, p<0.0001. (B,D) Latency was measured in bulk splenocytes by determining the frequency of viral genome-positive splenocytes by limiting dilution PCR. Briefly, intact splenocytes from mice at 16 dpi (B) or 42 dpi (D) were serially diluted and subjected to nested PCR reactions to detect viral ORF50. (C,E) Reactivation was measured by coculturing serial dilutions of intact splenocytes (C) or peritoneal exudate cells (E) from infected mice 16 dpi onto a monolayer of MEFs and scoring for CPE 2-3 weeks post-coculture. For the limiting dilution analyses, curve fit lines were determined by nonlinear regression analysis. Using Poisson analysis, the intersection of the nonlinear regression curves with the dashed line at 63.2% was used to determine the frequency of cells that were either positive for the viral genome (B,D) or reactivation (C,E). Data are generated from two (D) or three (B,C,E) independent experiments with 4 to 5 mice per group for IKKα-SA1 and IKKα-SA.STOP1 viruses. Error bars indicate SEM. Significance was determined by unpaired two-tailed t-test; *, p=0.01.

Macrophages of the peritoneal compartment comprise another reservoir of MHV68 latency (33). Peritoneal exudate cells isolated 16 days post-intraperitoneal infection revealed nearly identical levels of reactivation from latency in mice infected with IKKα-SA1 (1/2,218) or IKKα-SA.STOP1 (1/1,614) **(FIG 4E)**. Taken together, upon direct administration of the IKKα-SA1 mutant to the peritoneal cavity, latency was disrupted in splenocytes but not peritoneal exudate cells at 16 dpi. NIK-dependent non-canonical IKKα signaling has route-and tissue-dependent roles in supporting gammaherpesvirus latency *in vivo*.

## DISCUSSION

The MHV68 pathogen-host system provides a tractable model of gammaherpesvirus infection and latency establishment, enabling the contribution of host factors to specific tissues and stages of infection to be determined. This is the first investigation of the role that IKKα, an important mediator of B cell signaling, plays in the productive replication of MHV68 *in vitro* and the establishment and maintenance of latency *in vivo* in mice with an intact B cell reservoir. We engineered a recombinant virus that expresses a dominant negative form of IKKα, named IKKα-SA, with S176A and S180A mutations that prevent phosphorylation by the upstream NIK kinase. We demonstrate that IKKα-SA virus mediated repression of the non-canonical NF-κB pathway through impairment of LTβR-mediated activation of the NF-κB p52 subunit in primary fibroblasts **(FIG 1C)**. This inhibition did not result in a replication defect in primary fibroblasts in culture or in the lungs of mice following intranasal inoculation **(FIG 2B)**. Although early infection in the lung was unaffected, the IKKα-SA mutant was severely defective in colonization of the spleen and in the establishment of latency as compared to the IKKα-SA.STOP control and WT MHV68 at 16 dpi **(FIG 3A-3B)**. Concordantly, reactivation was undetectable in splenocytes infected with the IKKα-SA mutant **(FIG 3C)**. Even colonization of the spleen after intraperitoneal inoculation, a more permissive route of MHV68 infection, was moderately impaired **(FIG 4B)**. In striking contrast, virus latency and reactivation within the peritoneal compartment, comprised primarily of macrophages, was not impacted by IKKα-SA infection after intraperitoneal inoculation **(FIG 4E)**. Taken together, non-canonical NF-κB signaling is not required for lytic infection, but functions as a critical host determinant of latency in the B cell compartment.

For MHV68 to establish a long-term infection after intranasal inoculation, it must productively infect multiple lineages of cells from the initial sites of mucosal tissue infection to dissemination to lymph nodes and infection of B cells that drive hematogenous dissemination (10). In our model, we observe a specific reduction in latency establishment in the lymphoid compartment **(FIG 3B)**. Although we engineered our construct to co-express an IRES-driven H2b-YFP transgene, we found it was not expressed consistently in latently infected cells and thus we could not elucidate how blockade of non-canonical signaling affected trafficking to the spleen using flow-based cytometric profiling. To identify cell-type specific blockades in infection, future studies will move the IKKα-SA transgene to a virus with a different reporter construct to better enable the virus to be tracked as it transits the lymph nodes and disseminates to the spleen.

The gammaherpesviruses infect multiple cell types but preferentially establish latency in B cells, suggesting their fate is determined by interaction with the B cell signaling milieu. Non-canonical NF-κB signaling is highly active at multiple stages of B cell development, including entry into the germinal center (13, 30, 31, 34-37). The non-canonical pathway is activated by a set of receptors that are expressed primarily on B cells (37), such as the BAFFR. Impairing IKKα, a critical kinase in this signaling cascade, in the bone marrow compartment leads to severe maturation defects in early maturation and subsequent survival of the germinal center (30, 31). Even impairment of upstream non-canonical signaling in mice lacking the BAFF receptor manifests in severely reduced mature B cell populations and formation of only transient germinal centers (35, 37). Disruption of the non-canonical NF-κB signaling pathway through genetic knockouts has revealed some contributions of this pathway to gammaherpesvirus pathogenesis. MHV68 infection of BAFF receptor knock-out mice results in reduced latency establishment, albeit in the context of reduced B cell reservoirs (26). Likewise, impairment of CD40, a critical component of B cell survival and maturation (38, 39), including in the germinal center, reduced MHV68 latency. In bone marrow chimeric mice generated from CD40+/+ and CD40-/- donors, latency was established and maintained in long-term CD40+ B cells that could transit the germinal center. CD40- B cells, on the other hand, could be infected, but latency was progressively lost. Genetic deletion of Lymphotoxin α, a non-canonical signaling receptor critical for splenic architecture development but not for B cell survival or function (40), had slightly enhanced lytic replication, but no defect in latency establishment (41). By examining infection in the context of a normal B cell compartment, our model uncovers that functional non-canonical NF-κB signaling is a necessary determinant in driving the establishment of latency after intranasal infection (**FIG 3B)**. Infection via peritoneal inoculation, a more permissive and direct route of infection, was not sufficient to restore the establishment of latency **(FIG 4B)**, further highlighting the importance of IKKα signaling in latency establishment. The observation of latent infection in transitional B cells (42), an immature B cell subset that requires non-canonical NF-κB for survival, suggests that this pathway may also contribute to the establishment of new reservoirs and maintenance of long-term latency. Taken together, these findings help to underscore an interaction of gammaherpesvirus signaling with this core B cell pathway in the establishment of life-long infection.

While non-canonical pathway inhibition negatively affected latency, we found that MHV68 did not require IKKα for lytic replication based on comparable levels of virus production in infected fibroblast cells expressing the IKKα-SA transgene **(FIG 2)** and in cells depleted for IKKα using a tamoxifen-inducible Cre recombinase **(FIG S3)**. Reactivation from latency also appeared unimpaired after intraperitoneal infection; albeit reactivation scored lower for IKKα-SA than with IKKα-SA.STOP virus (**FIG 4C)**, the decrease in reactivation was consistent with the initial defect in latency establishment (**FIG 4B)**. Canonical NF-κB activation, which is more broadly activated than the non-canonical pathway, has been shown to play a deleterious role in lytic replication of MHV68 (43) and in lytic reactivation of EBV and KSHV from latency (44, 45). Our lab reported that NF-κB signaling influences occupancy by the major viral gene transactivator, RTA, at the origin of lytic replication in latent B cells, including those of infected mice (46). The gammaherpesviruses have strategies to repress canonical NF-κB activation by targeting the NF-κB subunit p65 for degradation (47-49). Given the importance of ubiquitin moieties in modulating TRAF2, TRAF3, and cIAP1/2 activity (50-53) and mediating NIK accumulation and p100 degradation, non-canonical pathway signaling is a potential target for viral modulation at the level of ubiquitination. The tegument protein ORF64 functions as a viral deubiquitinase (54) and plays an important role in innate immune evasion that is dependent on its ubiquitinase activity (55), but the ubiquitin modification states of non-canonical pathway signaling intermediates were not examined. There has been no systemic screen for gammaherpesvirus regulators of the non-canonical NF-κB pathway as reported for canonical NF-κB (19). Additional studies that more closely examine the activation status of both NF-κB pathways in cell types of physiologic relevance to gammaherpesvirus pathogenesis would better define the specific conditions that drive latency over deactivation of these pathways that might influence an abortive infection or lytic fate.

The oncogenic gammaherpesviruses activate NF-κB signaling upon infection (56, 57) and this activation promotes cell survival (58, 59). Several key modulators have been identified. Notably, LMP1 and vFLIP, two major latent proteins of EBV and KSHV, activate both the canonical and non-canonical NF-κB pathways and indicate that each signaling pathway is critical for B cell latency. EBV LMP1 mimics a constitutive activation of tumor necrosis factor receptors, as its carboxy-terminal activating region 1 (CTAR1) recruits TRAF2 and TRAF3, activates NIK and IKKα (60). Deletion of the CTAR1 sequence from the EBV genome significantly impairs the ability of EBV to transform B cells (61). While multiple signaling pathways are triggered by CTAR1 (62), CRISPR-based screening identified an essential role for p52 in the survival and growth of EBV transformed lymphoblastoid cell lines (63), confirming the critical function of IKKα in the maintenance of latency. KSHV vFLIP binds and activates IKKα and triggers non-canonical NF-κB signaling in a NIK-independent manner; vFLIP also increases the expression level of p100 (64, 65). Deletion of vFLIP results in apoptosis of infected B cells and lytic reactivation (66). Thus, activation of NF-κB is a striking feature of EBV and KSHV latently infected cells that underlies oncogenic processes. Direct counterparts to these oncogenes are not found in MHV68; MHV68 may rely on the microenvironment for these cues or perhaps a viral modulator remains to be discovered.

Our findings indicate that the non-canonical NF-κB signaling pathway is essential for the establishment of latency in the lymphoid tissue of mice infected with the murine gammaherpesvirus pathogen MHV68, recapitulating the roles for this signaling pathway described in the human gammaherpesviruses. We have previously reported that interference with canonical signaling disrupts virus-driven pathology (67). Disrupting the signals that govern latency, perhaps by antagonizing cytokines that drive NIK activation, may provide a novel and effective therapeutic avenue to combat gammaherpesvirus latency and lymphoproliferation.

## MATERIALS AND METHODS

### Mice and ethics statement

Wild-type C57BL/6 mice were purchased from Harlan/Envigo RMS (Indianapolis, IN) or Jackson Laboratories (Bar Harbor, ME). All animal protocols were approved by the Institutional Animal Care and Use Committee of Stony Brook University.

### Cell culture

Primary and immortalized C57BL/6 murine embryonic fibroblasts (MEFs) were harvested and maintained in Dulbecco’s modified Eagle medium (DMEM) supplemented with 10% fetal bovine serum (FBS), 100 U/ml of penicillin, 100 mg/ml of streptomycin, and 2 mM L-glutamine (10% DMEM) at 37°C in 5% CO2. Primary MEFs at passages 2 and 3 were used for viral growth curves and limiting dilution reactivation assays. Immortalized MEFS were passaged 1:3 every other day. NIH 3T12 murine fibroblast cells (ATCC CCL-164) were maintained in DMEM with 8% FBS, 100 U/ml of penicillin, 100 mg/ml of streptomycin, and 2 mM L-glutamine at 37°C in 5% CO_2_. Non-canonical pathway activation in MEFs was stimulated with 0.3μg/ml αLTβR clone 5G11 (Abcam, Cambridge, MA, USA) for 5 h.

### Viruses

MHV68-IKKαSA and MHV68-IKKαSTOP were generated using BAC-mediated recombination onto the WT MHV68 BAC (68, 69). First, the ECMV IRES was cloned out of pIRES (Clontech, Mountain View, CA, USA) into pEYFP-N1 30bp upstream of the eYFP. The pCMVIE-IRES-eYFP-SV40polyA sequence was excised using restriction digestion, blunted using the Klenow fragment of DNA polymerase (NEB, Ipswitch, MA, USA), and blunt-end ligated into the TOPO-TA blunt vector (Thermo-Fisher Scientific, Waltham, MA, USA). Human IKKαSA was amplified from pBIP-IKKαSA (70) and ligated into the TOPO-TA vector upstream of the IRES. The complete pCMVIE-IKKαSA-IRES-YFP-SV40polyA was excised using NsiI PstI double restriction digestion, gel purified, and ligated into NsiI and PstI sites in pUC19 (NEB, Ipswitch, MA, USA). A kanamycin resistance cassette and I-Sce cut site was cloned into the IRES of our construct. The entire construct was amplified by PCR, transformed into *E. coli* containing the WT BAC, and carbenicillin/kanamycin resistant clones were selected. I-Sce expression was induced at 37°C. Carbenicillin-resistant and kanamycin sensitive colonies were selected. BAC clones were sequenced using Illumina miSeq whole genome sequencing and assembled to the predicted genome sequence.

Next, this BAC was further modified to repair mutations in the IKKα coding sequence and modify the IRES and YFP reporter gene to improve poor YFP expression. First, a gBlock (Integrated DNA Technologies, Coralville, IA, USA) was synthesized with an extended IRES, a sequence deletion between the IRES and eYFP, and an H2B coding sequence fused to YFP in addition to a kanamycin-resistance cassette and an I-Sce cut site. Mutants were generated by *en passant* recombination onto the IKKα-SA BAC to generate the IKKα-SA-IRES-H2BYFP BAC (referred to as IKKα-SA). Next IKKα mutations were repaired by amplification of the IKKα gene from the BAC, site directed mutagenesis, followed by *en passant* recombination as described above. Next, to generate an IKKαSTOP virus, all-frames STOP cassettes were inserted after nt 111 (5’-<u>GTGATTAGCTGA</u>GGCGGCCGC-3’) and 1309 (5’-TTAATTAA<u>GTGATTAGCTGA</u>-3’) of the IKKα-SA gene using sequential site-directed mutagenesis of PCR products from the IKKα-SA BAC followed by *en passant* recombination as described above. Intended mutations in IKKα (IKKα-SA) and insertion of STOP cassettes (IKKα-SA.STOP) were confirmed by Sanger and whole genome sequencing.

Two independent recombinant MHV68 BACs were derived for IKKα-SA and for IKKα-SA.STOP, followed by transfection and passaging through Vero-CRE cells to remove the loxP-flanked BAC sequence and then final production in NIH3T12 cells. Virus stocks were concentrated to > 1 × 10^8^ PFU/ml by centrifugation at 4°C for 120 min at 13,000 x g in a Dupont GSA rotor (Wilmington, DE).

### Sequence analysis

To validate the sequences of our mutant viruses, BAC DNA was prepared using Qiagen columns (Germantown, MD), and a DNA library was prepared using Illumina Nextera DNA Library Preparation kit (Illumina, San Diego, CA). Fragmented and tagged DNA was subjected to MiSeq at the Stony Brook Microarray Facility. Whole-genome sequencing analysis was performed as paired-end DNA-seq; reads were imported into (Geneious v10.1.2) and trimmed using BBDuk (v36.92) and mapped to the MHV68-H2BYFP genome with medium sensitivity. Variants were detected with the parameters of a minimum depth coverage of 500 reads and minimum variant frequency of 0.4.

### Immunoblotting

Infected cells were lysed in RIPA buffer and quantified by Bradford assay (BioRad, Berkeley, CA) and protein was diluted RIPA Buffer (150 mM sodium chloride, 1.0% IGEPAL CA-630, 0.5% sodium deoxycholate, 0.1% sodium dodecyl sulfate, 50 mM Tris pH 8.0) supplemented with a protease inhibitor cocktail (Sigma, St. Louis MO) and PMSF before boiling at 95° C for 5 min. Proteins were separated on 10% SDS-PAGE and transferred to polyvinylidene fluoride membrane. Primary antibodies against IKKα (Cell Signaling Technology, Rabbit polyclonal, Danvers, MA, USA) and ORF57 (kindly provided by Dr. Paul Ling, Baylor College of Medicine) (71) were used. Detection was performed with HRP-conjugated anti-rabbit IgG or anti-mouse IgG (GE Healthcare, Buckinghamshire, UK). Data was captured by GE CCD camera and analyzed by ImageQuant software (v7.0, GE Healthcare).

### NF-κB p52 ELISA

Stimulated MEFs were washed in cold PBS and scraped into 1 ml cold PBS. Cells were pelleted by centrifugation at 600xg and resuspended in 300-500 μl of hypotonic lysis buffer (10 mM HEPES pH 7.9, 10 mM KCl, 1.5 mM MgCl_2_, 0.1 mM EDTA, 1 mM DTT, 0.5 mM PMSF, protease inhibitor cocktail (Sigma, St. Louis MO)). Cells were incubated for 15 min on ice, then 5% volume of 10% NP-40 was added to cells. Cells were vortexed for 15 s then centrifuged at 10,000 xg for 10 min. Supernatant was stored as the cytoplasmic fraction. The remaining pellet was washed twice in 500 μl of hypotonic lysis buffer, then resuspended in 50 μl of complete lysis buffer (Active Motif, Carlsbad, CA, USA). Pellets were shaken at 4°C for 2 h, then centrifuged at 10,000 xg for 10 min. The supernatant was saved as the nuclear fraction. The Active Motif TransAM p52 ELISA kit was used to quantify p52 activation in 20 μg of nuclear extract, according to manufacturer’s instructions.

### In vitro infections

Immortalized C57BL/6 murine embryonic fibroblasts (MEFs) were plated at a density of 0.9 × 10^5^ cells/mL one day prior to infection. Infection was carried out at multiplicity of infection (MOI) of 0.05 in a low-volume for 1 hr at 37°C, with rocking back-and-forth every 15 mins, followed by the addition of fresh complete media and incubated at 37°C. Cells were harvested at the indicated time points and freeze-thawed 5 times.

### Plaque assays

NIH 3T12 cells were plated at a density of 0.9 × 10^5^ cells/mL one day prior to infection. Serial dilutions of cell homogenate were added to the cell monolayer for 1 hr at 37°C, with rocking every 15 minutes, followed by an overlay of 5% methylcellulose (Sigma) in cMEM and incubated at 37°C. After 7-8 days, cells were fixed with 100% methanol (Sigma) and stained with 0.1% crystal violet (Sigma) in 20% methanol, and plaques were scored.

### Viral pathogenesis assays

For intranasal infection, mice were lightly anesthetized using isoflurane and infected with 1,000 plaque forming units (pfu) of virus in a 20 μl bolus of 10% cMEM applied to the nose. For intraperitoneal infection, mice were lightly anesthetized using isoflurane and administered with 1,000 plaque forming units (pfu) of virus in 500 μl of 10% cMEM in the peritoneal cavity.

For acute titers, mice were euthanized with isoflurane at the indicated days post infection, and the left and right lungs were removed and frozen at ^-^80°C. Lungs were disrupted in 1 ml of 8% cMEM using 1 mm zirconia beads in a bead beater (Biospec, Bartlesville, OK) and analyzed by plaque assays as described above.

To analyze latently infected cells, mice were euthanized with isoflurane at 16 or 42 dpi. Spleens were excised, homogenized, and resuspended in 10% cMEM. Peritoneal exudate cells were isolated by peritoneal injection of 10 ml of 10% cMEM, followed by agitation of the abdomen and withdrawal of the peritoneal wash by syringe. For quantitation of latency, limiting-dilution nested PCR with primers for the MHV68 ORF50 region was used to determine the frequency of virally infected cells as previously described (72). Briefly, frozen samples were thawed, resuspended in isotonic buffer, counted, and plated in serial three-fold dilutions in a background of 10^4^ NIH 3T12 murine fibroblasts into a 96-well plate. The resultant PCR products were resolved in 2% agarose gels and each dilution was scored for amplimer of the expected sizes. Control wells containing uninfected cells or 10, 1, and 0.1 plasmid copies of ORF50 target sequence were run with each plate to ensure single-copy sensitivity and no false positives.

For quantitation of reactivation, a limiting-dilution reactivation assay was performed as previously described (72). Briefly, bulk splenocytes in 10% cMEM were plated in serial 2-fold dilutions (starting with 10^5^ cells) onto MEF monolayers in each well of a 96-well tissue culture plate. Twelve dilutions were plated per sample, and 24 wells were plated per dilution. Wells were scored for cytopathic effect at 14 and 21 d after plating. To detect preformed infectious virus, parallel samples of mechanically disrupted cells were plated onto MEF monolayers.

### Statistical Analyses

All data were analyzed using GraphPad Prism software (GraphPad Software, http://www.graphpad.com, La Jolla, CA). Titer data were analyzed with unpaired t-test or one-way ANOVA for multiple groups. Growth curve data were analyzed with two-way ANOVA with Tukey’s multiple comparison test. Based on the Poisson distribution, the frequencies of reactivation and viral genome–positive cells were obtained from the nonlinear regression fit of the data where the regression line intersected 63.2%. Extrapolations were used for samples that did not intersect 63.2%. The log_10_-transformed frequencies of genome-positive cells and reactivation were analyzed by unpaired, two-tailed t-test or one-way ANOVA for multiple groups.

## ACKNOWLEDGEMENTS

We thank Hong Wang for technical assistance with library preparation for MiSeq; Sumar Hayan and Steven Reddy for technical support; and Jean Rooney, Nicole Motta, and Laurie Levine for their assistance with mouse techniques and animal husbandry. Special thanks to Drs. James Bliska, Nicholas Carpino, Adrianus van der Velden, Douglas White and all members of the Krug laboratory for helpful discussions.

B.C. was supported by NIAID T32AI007539, V.K., Q.D., and L.T.K. were supported by an American Cancer Society Research Scholar Grant RSG-11-160-01-MPC. D.G.O. was supported by the Gundersen Medical Foundation. This research was supported in part by the Intramural Research Program of the NIH, Center for Cancer Research.

B.C., V.K., I.D., Q.D., D.G.O., J.A.B., K.B.M., and L.T.K. and L.T.K., designed the experiments. B.C., V.K., I.D., Q.D., D.G.O., J.A.B., and L.T.K. executed the experiments. B.C., X.L., V.K., I.D., Q.D., D.G.O., J.A.B., and L.T.K. analyzed the data. B.C., X.L., and L.T.K. prepared the manuscript.

