## Supplemental Materials for "IKKalpha-Mediated Non-canonical NF-kappaB Signaling is Required to Support Murine Gammaherpesvirus 68 Latency *In Vivo*"

**SUPPLEMENTAL MATERIAL**

### Supplemental Methods

**Viral genome analysis.** For restriction fragment length polymorphism (RFLP) analysis, BAC DNA was digested using HindIII, EcoRI, or BamHI and resolved on a 2% agarose gel at low voltage for 8 hrs.

**Immunoblotting.** Stimulated cells were lysed in RIPA buffer and quantified by Bradford assay (BioRad, Berkeley, CA) and protein was diluted RIPA Buffer (150 mM sodium chloride, 1.0% IGEPAL CA-630, 0.5% sodium deoxycholate, 0.1% sodium dodecyl sulfate, 50 mM Tris pH 8.0) supplemented with a protease inhibitor cocktail (Sigma, St. Louis MO) and PMSF) before boiling at 95$^{o}$ C for 5 min. Proteins were separated on 10% SDS-PAGE and transferred to polyvinylidene fluoride membrane. Primary antibodies against p100 (Cell Signaling Technology, Rabbit polyclonal), IKKα (Cell Signaling Technology, Rabbit polyclonal, Danvers, MA, USA), p65 (Cell Signaling Technology, Clone C22B4), GAPDH (Sigma-Aldrich, Rabbit Polyclonal), and Lamin A/C (Cell Signaling Technology, Clone 4C11) were used. Detection was performed with HRP-conjugated anti-rabbit IgG or anti-mouse IgG (GE Healthcare, Buckinghamshire, UK), Data was captured by GE CCD camera and analyzed by ImageQuant software (v7.0, GE Healthcare).

**IKKα deletion in MEFs.** Mice bearing germline $CreER^{T2}/IKK\alpha^{fl/fl}$ on a C57BL/6 background were bred in the Stony Brook University facility (1). MEFs generated from $CreER^{T2}/IKK\alpha^{fl/fl}$ mice were treated daily with 100 μM tamoxifen (Sigma-Aldrich, St. Louis, MO, USA) + 10% CMEM for 8 days prior to infection.

NF-kappa B kinases alpha and beta are both essential for high mobility group box 1-mediated chemotaxis. J Immunol 184:4497-509.

### Supplemental Figure Legends

**FIG S1.** Analysis for restriction fragment length polymorphisms in recombinant MHV68 BACs. (A) Schematic of IKKα-SA and IKKα-SA.STOP viruses with restriction sites and resulting band sizes. B: BamHI, E: EcoRI, H: HindIII. (B) Recombinant viral BACs were analyzed by restriction digest. In the BamHI digest, IKKα-SA virus has the appearance of two bands 920 bp and 1,381 bp, and IKKα-SA.STOP virus has the appearance of two bands 940 bp and 1,402 bp (red arrows), all compared to WT BAC- parental. The EcoRI digest reveals the appearance of a 5,438 bp band and 5,615 bp band in both IKKα-SA and IKKα-SA.STOP viruses compared to WT BAC- parental. Lastly, in the HindIII digest, there is the appearance of a 6,570 bp band in both IKKα-SA and IKKα-SA.STOP compared to WT BAC- parental.

**FIG S2.** αLTβR stimulation induces p100 degradation in MEFs. (A) Primary WT MEFS were stimulated with 1 μg/ml αLTβR for 18 h or 50 ng/ml TNFα for 15 min. Cell lysates were fractionated to collect cytoplasmic and nuclear fractions. p100 cleavage and p65 were detected by immunoblotting. Lamin A/C, control for nuclear fraction; β-tubulin, control for cytoplasmic fraction.

**FIG S3. IKKα signaling is not necessary for MHV68 lytic infection.** MEFs were generated from AFTC ($CreER^{T2}/IKK\alpha^{fl/fl}$) mice and treated with tamoxifen daily for 8 days to drive IKKα deletion ($IKK\alpha^{-/-}$). (A) Cells were lysed and probed for IKKα protein to verify loss. GAPDH was used as a loading control. (B) $IKK\alpha^{-/-}$ and WT MEFs were infected with MHV68-WT at an MOI of 5. Cells were harvested at 24 and 48 hpi, and virus replication was measured by plaque assay. Timepoints were measured in triplicate. Data was analyzed by unpaired t-test, n.s.=$p>0.05$.

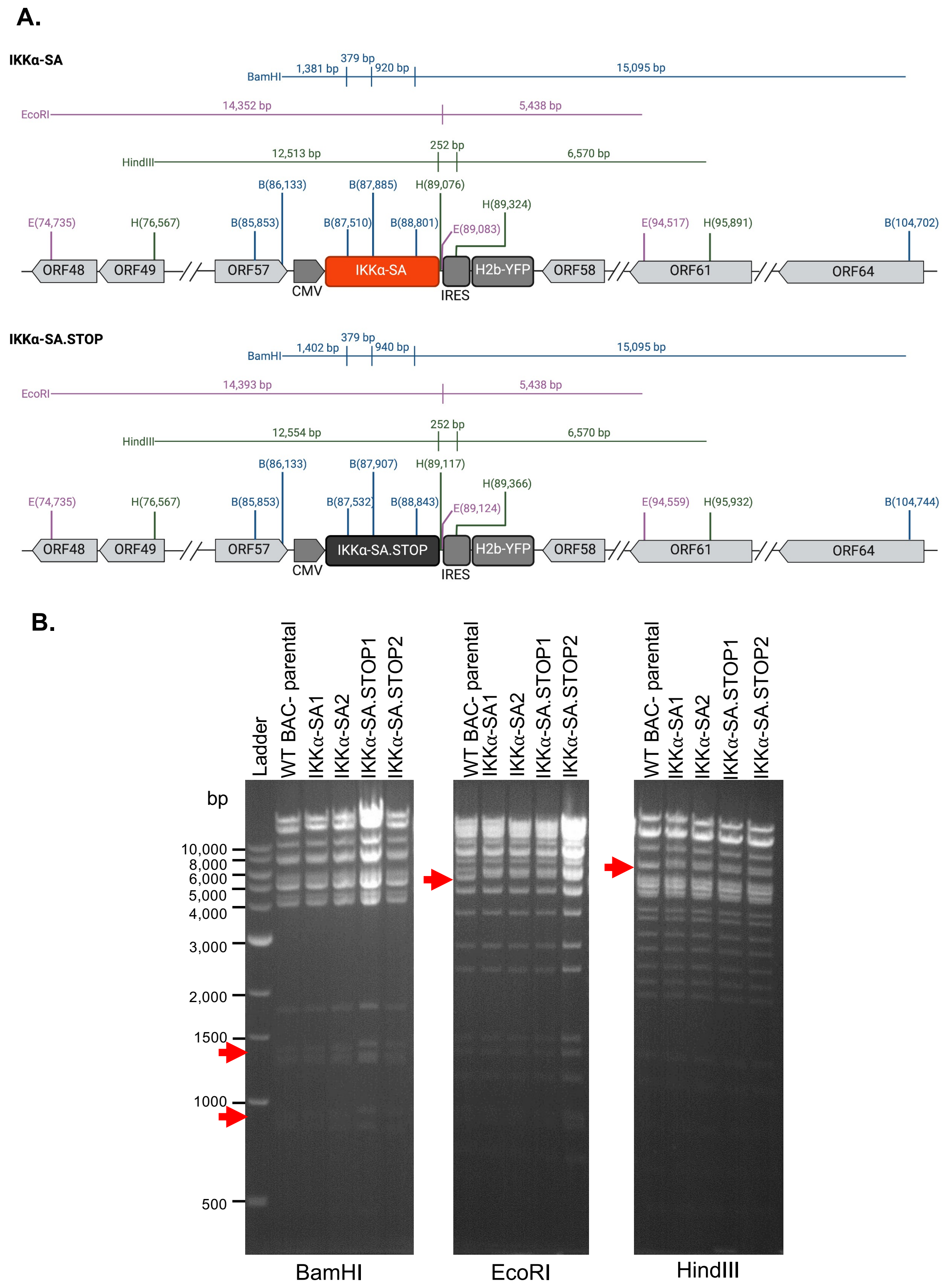


**FIG S1.** Analysis for restriction fragment length polymorphisms in recombinant MHV68 BACs. (A) Schematic of IKKα-SA and IKKα-SA.STOP viruses with restriction sites and resulting band sizes. B: BamHI, E: EcoRI, H: HindIII. (B) Recombinant viral BACs were analyzed by restriction digest. In the BamHI digest, IKKα-SA virus has the appearance of two bands 920 bp and 1,381 bp, and IKKα-SA.STOP virus has the appearance of two bands 940 bp and 1,402 bp (red arrows), all compared to WT BAC- parental. The EcoRI digest reveals the appearance of a 5,438 bp band and 5,615 bp band in both IKKα-SA and IKKα-SA.STOP viruses compared to WT BAC- parental. Lastly, in the HindIII digest, there is the appearance of a 6,570 bp band in both IKKα-SA and IKKα-SA.STOP compared to WT BAC- parental.

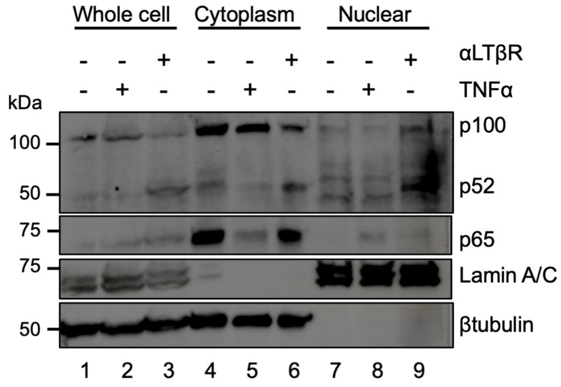


**FIG S2.** αLTβR stimulation induces p100 degradation in MEFs. (A) Primary WT MEFS were stimulated with 1 μg/ml αLTβR for 18 h or 50 ng/ml TNFα for 15 min. Cell lysates were fractionated to collect cytoplasmic and nuclear fractions. p100 cleavage and p65 were detected by immunoblotting. Lamin A/C, control for nuclear fraction; β-tubulin, control for cytoplasmic fraction.

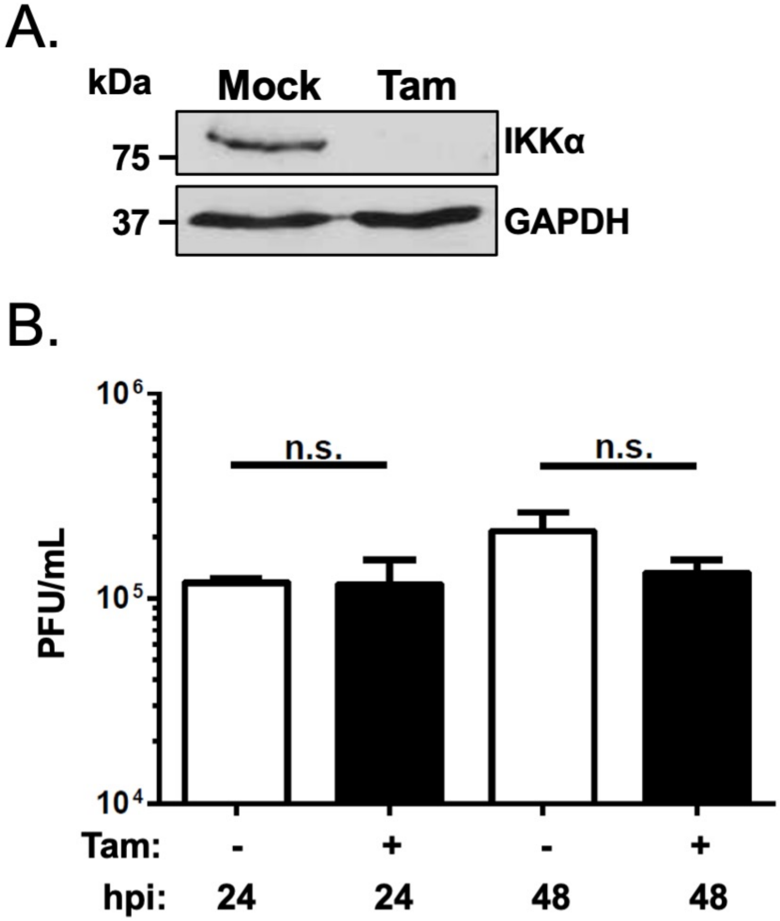


**FIG S3. IKKα signaling is not necessary for MHV68 lytic infection.** MEFs were generated from AFTC (*CreER*$^{T2}$/*IKKα*$^{fl/fl}$) mice and treated with tamoxifen daily for 8 days to drive IKKα deletion. (A) Cells were lysed and probed for IKKα protein to verify loss. GAPDH was used as a loading control. (B) IKKα$^{-/-}$ and WT MEFs were infected with MHV68-WT at an MOI of 5. At 24 and 48 hpi cells were harvested and viral replication was measured by plaque assay. Timepoints were measured in triplicate. Data was analyzed by unpaired T test, n.s.=$p>0.05$.
